# In-situ metagenomics: A platform for rapid sequencing and analysis of metagenomes in less than one day

**DOI:** 10.1101/2023.01.25.525498

**Authors:** Javier Tamames, Diego Jiménez, Álvaro Redondo, Sandra Martínez-García, Asunción de los Rios

## Abstract

We present here a complete system for metagenomic analysis that allows performing the sequencing and analysis of a medium-size metagenome in less than one day. This unprecedented development was possible due to the conjunction of state-of-the art experimental and computational advances: a portable laboratory suitable for DNA extraction and sequencing with nanopore technology; the powerful metagenomic analysis pipeline SqueezeMeta, capable to provide a complete analysis in a few hours and using scarce computational resources; and tools for the automatic inspection of the results via a graphical user interface, that can be coupled to a web server to allow remote visualization of data (SQMtools and SQMxplore). We have tested the feasibility of our approach in the sequencing of the microbiota associated to volcanic rocks in La Palma, Canary Islands. Also, we did a two-day sampling campaign of marine waters in which the results obtained the first day guided the experimental design of the second day. This system offers the possibility of prospecting any habitat in a quick way, obtaining both taxonomic and functional information about the target microbiome.

## Introduction

The popularization of portable sequencers, especially those based on nanopore technologies (Deamer et al., 2016), has created the possibility of having rapid sequencing data which can be very valuable in several contexts, for instance in clinical scenarios of disease control or epidemics (Quick et al., 2015, 2016). Also, the portability of these devices has been explored *in situ*, for example in oceanographic expeditions or in the Antarctic ice (Gowers et al., 2019; Johnson et al., 2017; Lim et al., 2014), illustrating the capability of producing sequences readily. This allows to envision the capacity of designing dynamic sampling campaigns, where the planning of the whole campaign can be driven by the results being produced. This can be important, for instance, whenever the sampling takes place in remote regions for which is desirable to have prompt data acquisition to prevent suboptimal results. It will be valuable also in any study where following the course of a microbiome in real time is necessary, for example when monitoring microbial blooms (Nowinski et al., 2019), assessing the quality of drinking waters (including security and bioterrorism) (Turingan et al., 2013), or controlling food processing issues like fermentations (De Filippis et al., 2017, Walsh et al., 2017). While standard amplification approaches (metabarcoding) can be useful in some of these cases (for instance, for detecting particular organisms in a sample), they may present issues related to biases in the amplification, and are usually limited to study taxonomic composition and/or very specific functions (Laudadio et al., 2018). When the objective is to obtain a full functional profile of the whole community, or we suspect that our sample may contain many interesting unknown organisms, metagenomics is a most sensible option (Zepeda el at, 2015). Metagenomics is a powerful tool for gaining insight on microbial communities, and has become a standard procedure for analyzing the structure and functionality of microbiomes.

The bottleneck of metagenomics is often the complexity of the associated bioinformatic analysis. To relieve this burden, we developed the SqueezeMeta pipeline (Tamames & Puente-Sánchez, 2019a) with several objectives in mind: 1) offering a fast and easy-to use platform for performing the complete analysis of metagenomes. Our goal was to include all the usual steps in metagenomic analysis with state-of-the-art tools, but making them attainable to all users, no matter their bionformatic skills. 2) Breaking the dependence on large computers, making it able to run with scarce resources, and even in laptops. And 3) Providing additional tools for performing the statistical analysis and sharing the results.

Since then, we and others have tested the ability of SqueezeMeta to fulfill these requirements in many different instances. These capabilities make SqueezeMeta an optimal system for analyzing metagenomic data in all settings, even under difficult environmental conditions, and with poor logistic setups and limited computational resources.

Lately, our challenge has been to be able to produce a complete metagenomic analysis in less that 24 hours, directly on the sampling spot, without relying on electrical power or internet connectivity. This will make our system capable to work in any circumstance and in any environment (including the most remote ones), and to obtain real-time results that can shared with others on-the-fly. To do so, we devised a platform composed of several different modules:

1. A portable DNA extraction laboratory, small enough to be carried by one person, to isolate DNA suitable for analysis.
2. a MinION nanopore sequencer for processing that DNA, producing metagenomic sequences.
3. The bioinformatic pipeline SqueezeMeta, running in a small laptop, to analyze the metagenomic DNA sequences, and:
4. The stand-alone statistical package SQMTools (Puente-Sánchez et al., 2020) to perform statistical analysis of the data, coupled to our new SQMxplore library (https://github.com/redondrio/SQMxplore) which allows creating interactive web pages and interfaces for openly sharing the results.

These steps are summarized in Figure 1. For testing the feasibility of in-situ sequencing and the dynamic design of campaigns, we performed two different sampling and sequencing experiments. The first aimed to set up the protocol under field conditions, sequencing the microbiota associated to volcanic rocks on La Palma island (Canary Islands, Spain). The second aimed to design a two-day campaign in which the results of the first day can be used to determine the objectives for the second one. For this purpose, we chose sampling marine planktonic communities in the Ria de Vigo (Spain).

**Figure 1:**
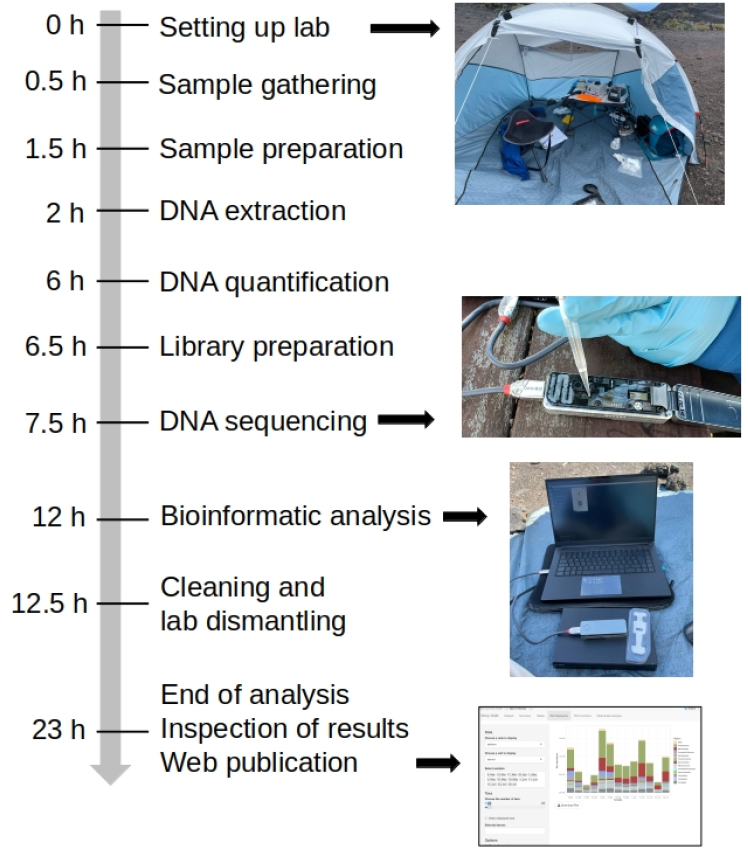
Approximate timeline of an in-situ metagenomic experiment. Time points in the left side are estimates, and refer to the starting time of the given step.

## Materials and Methods

### Portable DNA extraction laboratory

The portable laboratory is composed of the following items:

-MicroSpin centrifuge, yielding 12.500 RPM (ThermoFisher, Waltham, MA, USA)

-Table Vortex, lightened by removing the metal base (ThermoFisher, Waltham, MA, USA)

-Mini agate mortar and pestle, for homogenizing samples.

-MicroSpinner (ThermoFisher, Waltham, MA, USA)

-Two mini batteries to power up all systems (U’King Shenzhen Zhuo Qiong Technology Co., Ltd., China)

-PowerSoil DNA extraction kit (Weight: Qiagen NV, Venlo, Netherlands)

Optionally, in case of cold conditions, we can heat the devices using:

-3 Hand warmers (up to 60°C, Shenzhen Ziheng Technology Co., Ltd., China)

-2 portable thermal isolated containers

The extraction protocol implies bead-beating and centrifugation, that can be done efficiently using this portable equipment. Our tests of efficiency indicated that the yield is similar to that obtained with standard lab equipment (data not shown). Microbial DNA is usually scarce in environmental samples. Therefore, it is advisable to process several extraction tubes using the same filtration column, in order to collect as much DNA as possible. In our settings, we process 8 tubes per column. It is also advisable to perform a gentle bead-beating, in order to maintain DNA integrity as much as possible, which will be very important to obtain good results in the subsequent sequencing step. In addition, we have improved the results by purifying the extracted DNA using Omega Mag-Bind TotalPure NGS Beads (Omega Bio-Tek, Norcross, GA), which helps to preserve the life span of the flow cell.

All the devices are powered by a portable battery (222Wh/60000mAh) with autonomy for 12 hours of normal functioning. In case of cold conditions, we insulated the batteries and other equipment in an insulated lunch bag, heated by placing hand warmers in it. Cool conditions for storing some reagents are kept by using an insulated thermal container (portable 10 l camping fridge) with cold packs inside.

### Laboratory transportation and setting

All devices can be carried in a suitcase, or a medium backpack (60 liters). The total weight is around 13 Kg. A light camping tent is used to provide shelter and protection from sun, rain, moisture, or winds. Inside of the tent, a small folding table (1×1 meters) is sited as stable surface, together with a portable chair (Figure 2).

**Figure 2:**
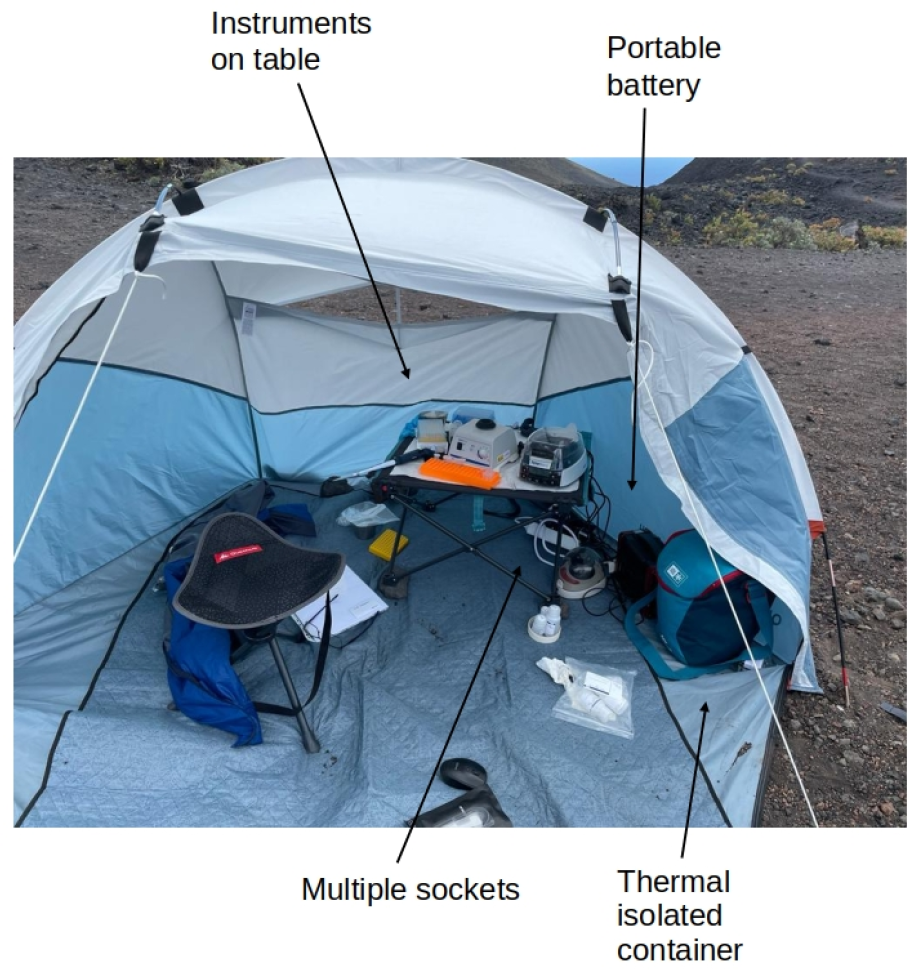
In-field setting of the portable laboratory

### DNA sequencing platform

The sequencing module is composed of the following items:

-Qubit 4 fluorometer (Invitrogen, ThermoFisher, Waltham, MA, USA)

-MinION sequencer (ONT, Oxford, UK)

-MinION flow cell (ONT, Oxford, UK)

-RAPid Sequencing Kit (ONT, Oxford, UK)

-Micro Thermocycler or portable water heater

-Laptop Schenker XMG Fusion 15 (16 Gb RAM, 8 core), with stand-alone MinKNOW software (v21.02., ONT, Oxford, UK)

First, the DNA concentration was measured using the Qubit fluorometer. This is needed to correctly adjust the amount of DNA to be introduced in the flow cell. The concentration of DNA obtained from environmental samples is variable, but can be rather low in lava rocks. Then, we calculated the volume of the DNA solution to be added for introducing 400 ng of DNA. As the volume of DNA extract for preparing the library is 10 ml, we estimated that a minimum DNA concentration of 40 ng/ml is needed. Several samples can be multiplexed in the same sequencing run.

The library is prepared using the RAPid kit from ONT, following manufacturer’s instructions, and barcoding the diverse samples with different tags. This kit includes a transposase that must be thermally inactivated. This can be done using a mini thermocycler, or simply heating water using a water immersion heater and incubating briefly the solution.

The sequencing time to reach the desired amount of sequence depends on several factors (DNA concentration, flow cell integrity, etc). In cold conditions, the MinION device and the laptop are protected by using insulated containers, which can be heated by placing hand warmers inside.

### Bioinformatics platform

The equipment needed for the bioinformatic analysis are the following:

-Same laptop than above (Schenker XMG Fusion 15), running the SqueezeMeta pipeline (https://github.com/jtamames/SqueezeMeta), R, the SQMTools, SQMxplore and Shiny R libraries installed. Internet connectivity is not needed for functioning, but of course would be necessary for sharing the data over the internet, if desired.

-Mini batteries to power up the laptop (U’King Shenzhen Zhuo Qiong Technology Co., Ltd., China)

SqueezeMeta is a fully automatic software that performs all the steps of the bioinformatic analysis of metagenomic data (Tamames & Puente-Sánchez, 2019a). The preferred mode of analysis implies assembling the raw sequences. But when the amount of sequencing is moderate, as in our case, the performance of the assembly decreases and it is advisable to run the analysis directly on the raw reads (Tamames et al, 2019b). Each read is then processed looking for ORFs and performing taxonomic and functional annotation for them, using the sqm_longreads program from the SqueezeMeta suite. The results are composed by a set of tables compiling all the information found for each read (including functional and taxonomic assignments), and statistics on the abundance of taxa and functions.

The drawback of using read annotation is that it consumes more resources and time, thus compromising our goal of performing the complete pipeline in less than 24 hours. Accordingly, the following strategy was used for the marine samples: Analyze the first three samples by co-assembly using an assembler such as Flye (Kolmogorov et al, 2019), Canu (Koren et al, 2017), or MEGAHIT (Li et al, 2015), to provide a quick analysis adequate to determine the most interesting spot for additional sampling. The first two are preferable, since they are optimized for working with MinION reads. The “--singletons” option of SqueezeMeta was used, allowing the addition the unassembled reads as new contigs. The second set of samples was analyzed using careful annotation of reads.

The analysis of the results is facilitated by the SQMtools R package (Puente-Sánchez et al, 2020), part of the SqueezeMeta suite. This library imports the tables resulting from the SqueezeMeta run and creates a R object that can be used to perform many different statistical analyses. SQMtools includes many prefabricated commands to obtain easily the most common types of plots and analyses.

The final step is the visualization and publication of results to make them accessible to the public. For this we use SQMxplore, which is a graphical user interface based on Shiny, a R library to build interactive web apps straight from R. SQMxplore takes the results from SqueezeMeta and SQMtools and displays them using a web browser. The data can be easily explored and shared, for instance by drawing histograms for the taxonomic composition of the sample, or the abundance of different functions. In this way, a remote user is able to access the results for inspection, without the need of (bio)informatic skills.

Taxonomic diagrams were plotted with Pavian (Breitwieser & Salzberg, 2020), via the sqm2pavian script of the SqueezeMeta pipeline. KEGG diagrams were done using the SQMtools interface to PathView (Luo & Brouwer, 2013).

### Sampling design: microbial communities on volcanic rocks

For the sequencing of microbiota associated to volcanic rocks, lava rock samples of two different ages were taken in May 2022 from lava fields in the south of La Palma island (Canary Island, Spain). Two main volcanic eruptions took place at the sampling site: San Antonio volcano (1677), and Teneguia volcano (1971). Each of them produced its own lava flows, which are very close and easily identifiable (Figure 3) (28°28’32"N 17°51’04"W).

**Figure 3:**
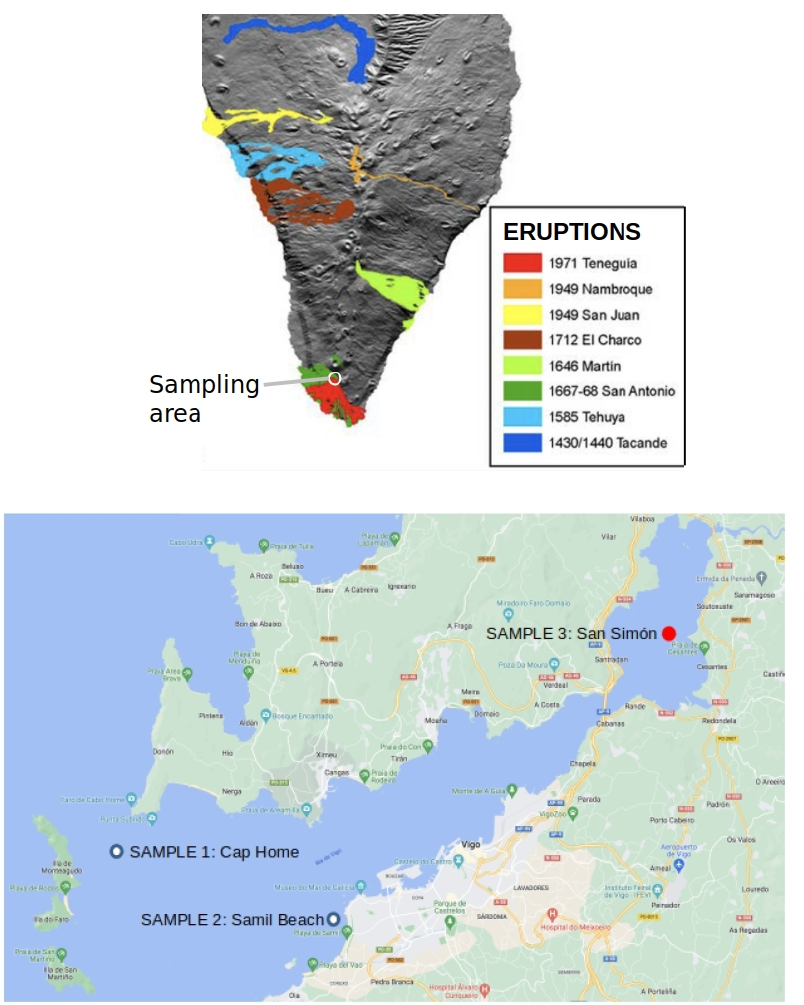
Upper: Recent eruptions in La Palma island, and location of the sampling spots in the confluence of Teneguia and San Antonio lava flows (Source: http://www.ign.es/resources/docs/IGNCnig/VLC-Teoria-Volcanologia.pdf). Lower: Sampling locations in Ria de Vigo

We took 5 subsamples (weighting approximately 100 grams each) in each of the spots and combined them to obtain one sample per sampling spot.,. We crumbled down them using a small mortar and pestle, to obtain a fine grained powder suitable for the PowerSoil extraction kit.

The resequencing of the volcanic samples for validation was performed using llumina NextSeq2000, in FISABIO (Valencia, Spain).

### Sampling design: planktonic microbial communities

We planned an oceanographic cruise in the Ría de Vigo (NW Iberian Peninsula). During the first day (July 12th 2022), surface (2 m) water samples were taken at three different locations: one in the outer part of the Ría, which is significantly influenced by oceanic waters (Cap Home, 42° 14.262’N 8° 52.325’W), one in the middle sector of the embayment in a anthropogenically affected area (Samil Beach, 42° 12.551’N 8° 46.983’W), and the last one in the inner part of the Ría with a relatively higher influence of riverine discharge (San Simón Bay, 42° 18.707’N 8° 37.926’W) (Figure 3). The three samples were processed and analyzed in order to choose the microbial community with the most interesting metagenomic profile, to repeat the sampling at the corresponding site the following day (July 14th 2022), increasing the sequencing depth of the analysis and the sampling resolution in the water column (2 m and 5 m).

Seawater samples were collected in 5 l acid-cleaned Niskin bottles and filtered through a 200 μm pore size mesh to remove larger zooplankton, in order to ensure good replication and facilitate filtration process. Subsequently, 12 l acid-washed polycarbonate bottles were gently filled and kept under dim light conditions, until arrival to the laboratory. Microplankton biomass was concentrated by means of sequential filtration through 3 and 0.2 μm pore-size polycarbonate filters at low vacuum pressure. Particles retained in the 3 μm pore-size filters were discarded, and microbial DNA was extracted from the 0.2 μm pore-size polycarbonate filters.

As explained above, approximately 5 liters of water were processed for each sample. When processing seawater samples, a first step of microplankton biomass concentration by means of vacuum filtration is needed. Onboard logistics did not allow to perform this filtration at sea, although this is a procedure often performed during oceanographic cruises. Therefore, water samples were taken to the laboratory at Estación de Ciencias Marina de Toralla (ECIMAT, Vigo) for filtration. The rest of the protocol remains unaltered.

## Data availability

SqueezeMeta and SQMtoos software are available at the following address: https://github.com/jtamames/SqueezeMeta.

SQMxplore software is available at: https://github.com/redondrio/SQMxplore Sequence data from volcanic rock and marine samples can be found at: https://saco.csic.es/index.php/s/s7tEaRLgL9wX3r8

## Results

### Volcanic samples

The goal of this experiment was to assess the differences in community structure in lava rocks of different ages (Teneguia and San Antonio samples), in order to shed light on the microbial colonization patterns of these rocks. Therefore, we were interested in determining the taxonomic profile of both samples.

We were able to reach the objective of completing the full protocol of sampling, DNA extraction, sequencing and in-situ analysis in less than 24 hours, powered all the equipment with batteries and in the absence of data connectivity. The amount of DNA obtained from these rocks was rather low: 14.7 ng/ml in Teneguía, and 32.1 ng/ml in San Antonio. In order to obtain a reasonable sequencing depth, the sequencing had to be extended for several hours, resulting in almost complete degradation of the flow cell. We sequenced a total of 286.4 Mb, 191 Mb for San Antonio and 95.4 Mb for Teneguia (Table 1). Raw reads for these samples were analyzed using the script sqm_longreads.pl from the SqueezeMeta pipeline (Table 1).

**Table 1:**
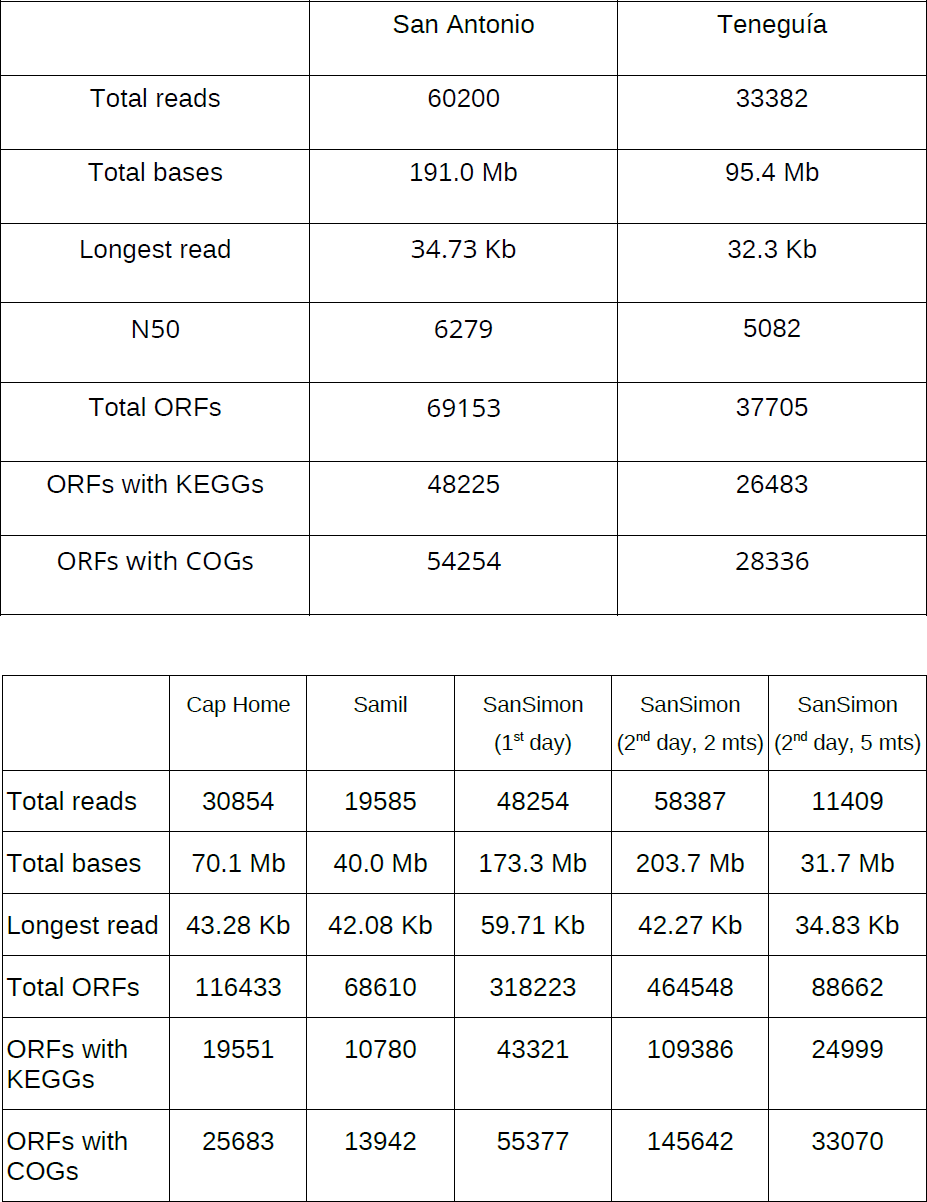
Sequencing and analysis data for both environments: Upper table: Volcanic rock samples. Lower table: Seawater samples

|  | San Antonio | Teneguía |
| --- | --- | --- |
| Total reads | 60200 | 33382 |
| Total bases | 191.0 Mb | 95.4 Mb |
| Longest read | 34.73 Kb | 32.3 Kb |
| N50 | 6279 | 5082 |
| Total ORFs | 69153 | 37705 |
| ORFs with KEGGs | 48225 | 26483 |
| ORFs with COGs | 54254 | 28336 |

Table 1: Sequencing and analysis data for both environments: Upper table:
|  | Cap Home | Samil | SanSimon<br>(1<sup>st</sup> day) | SanSimon<br>(2<sup>nd</sup> day, 2 mts) | SanSimon<br>(2<sup>nd</sup> day, 5 mts) |
| --- | --- | --- | --- | --- | --- |
| Total reads | 30854 | 19585 | 48254 | 58387 | 11409 |
| Total bases | 70.1 Mb | 40.0 Mb | 173.3 Mb | 203.7 Mb | 31.7 Mb |
| Longest read | 43.28 Kb | 42.08 Kb | 59.71 Kb | 42.27 Kb | 34.83 Kb |
| Total ORFs | 116433 | 68610 | 318223 | 464548 | 88662 |
| ORFs with KEGGs | 19551 | 10780 | 43321 | 109386 | 24999 |
| ORFs with COGs | 25683 | 13942 | 55377 | 145642 | 33070 |

The taxonomic profiles obtained by the analysis of the metagenomes can be seen in Figure 4. While the bacterial community structure is rather similar in both samples, marked differences were founf with respect to eukaryotic compounds. The composition of Ascomycota assigned to Lecanoromycetes (major class including lichen-forming fungi) differed between both samples. A clear predominance of sequences assigned to the genus *Letharia* (Lecanorales) and presence of *Cladonia* genus (Lecanorales) was observed in Teneguia lava rocks. However, in San Antonio samples, sequences assigned to the genera *Letharia* (Lecanorales) and *Lasallia* (Umbilicariales) were detected, but without the clear dominance of *Letharia* found in Teneguia samples. In addition, sequences assigned to the fungal orders Chaetotyriales and Leotiomycetes were only found in San Antonio samples. On the other hand, sequences assigned to *Trebouxia* (Chlorophyta, Trebouxiales), the most common photobiont of lichen-forming fungi, were also detected only in San Antonio samples. With respect of bacterial communities, differences in composition of the phyllum Actionabacteria were also found between both samples. These results reveal that the age of the lava mainly conditions the fungal composition and the establishment of lichen communities. Thus, it is demonstrated that this platform is useful to identify differences in microbial composition in the field, and focus subsequent sampling.

**Figure 4:**
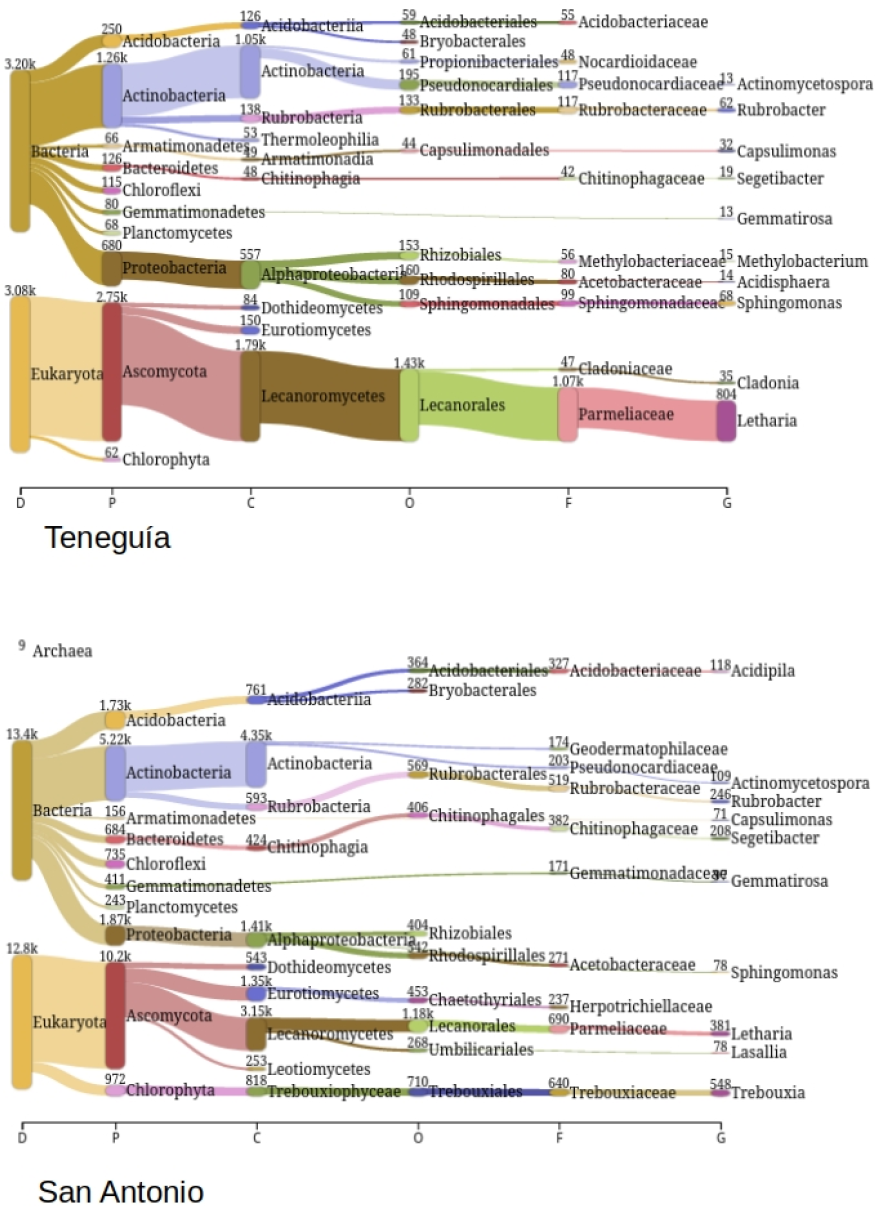
Taxonomic profiles of Teneguia and San Antonio metagenomic samples of lava rocks. Plots were done using Pavian (Breitwieser & Salzberg, 2020) and the sqm2pavian script of SqueezeMeta.

We also generated functional profiles for both samples, making it possible to analyze functional diversity exemplified by the abundance of genes involved in sulfur metabolism (Suppl Figure 1).

To validate our approach and demonstrate that it produces valid and usable results, we resequenced both samples using Illumina NextSeq2000, obtaining 20 million sequences per sample that were processed using the same SqueezeMeta pipeline than for MinION sequences. That is, analyzing the reads using the sqm_longreads.pl script. The results are shown in Suppl Figure 2, and indicate a very strong correlation between results from MinION and Illumina (In all cases, R^2^>0.94, p<0.01). Both taxa and functions abundances are very similar, with most abundant taxa and functions well preserved among them. Therefore, our in-situ MinION sequencing produces accurate results and can be used for studying functional and taxonomic composition of microbiomes.

### Marine water column samples

The objective of this experiment was to test the feasibility of planning a results-driven cruise, in which an initial sampling of different locations can serve to determine the most interesting spot to be further analyzed on subsequent days.

Our primary objective was to study sulfur metabolism in the Ria de Vigo. The Ria is characterized by high productivity due to upwelling events that promote the intrusion of nutrient-rich water to the embayment (Nogueira et al, 1997). This natural productivity and activities related to mussel farming are associated with an increased flow of organic matter to the seabed. Microbiological degradation of this organic matter consumes oxygen from the sediment interstitial water, promoting the development of anoxic zones where sulfate reduction and methane production processes coexist (García-Gil et al, 2003). We were interested in testing possible differences in some parts of the Ria, because sediment anoxic conditions have been shown to be more prevalent and shallower in the sediment cores from the inner part of the Ría (the San Simon Bay, which shows the characteristics of a typical estuary and is subjected to particularly important inputs of organic matter) compared to the middle or the outermost zones (which are subjected to oceanic influence). In fact, the highest sulfide concentrations are usually found in the inner zone of the Ría, the San Simon Bay (Ramírez-Pérez et al, 2015). A recent work (de Carlos et al, 2017) demonstrates important differences between the taxonomic composition of microbial communities living in shallow organic-rich estuarine sediments from San Simón Bay and in non-gassy sediments retrieved from the outer area of the ria. The authors suggest these differences are likely related to sediment type and differences in the cycling of organic matter, sulfur and methane (de Carlos et al, 2017).

The aim of the present work was to study the differences in microbial processes related to sulfur cycle in the water column in distinct sectors of the Ría de Vigo. Our hypothesis is that gas escapes from seafloor will differentially affect the sulfur cycling in the water column in distinct sectors of the Ría de Vigo. We decided to explore three locations of the Ria de Vigo, looking for the one with most interesting or most abundant genes related to sulfur metabolism. We performed two different samplings. During the first day (12^th^ July), we took microplankton surface samples in three different locations in the Ría de Vigo, sequenced DNA and analyzed the sequences in less than 24 hours. Metagenomic information recovered during the first day informed about sulfur metabolism in the three stations, and helped to choose the most interesting location to perform a more detailed analysis (increased vertical resolution of sampling and increased sequencing depth) during the second day.

After DNA extraction, we were able to retrieve the following DNA concentrations in the three spots: 8.60 ng/ml, 9.75 ng/ml in, and 21.8 ng/ml, for Cap Home, Samil Beach, and San Simón Bay samplings spots, respectively. These concentrations are below optimal, but still amenable to be sequenced.

Giving these concentrations, the three samples were barcoded and pooled using equimolar amounts of DNA. Subsequently, samples were put into the MinION flow cell for sequencing. To maximize flow cell survival, we decided to sequence for only 10 hours, as this was an exploratory analysis and consequently a large sequencing depth was not necessary.

We obtained 98.693 reads, corresponding to 283 Mb of sequence (Table 1). Even if we pooled equimolar amount of DNA for the tree samples, the result did not preserve equal quantities for each sample. Indeed, 48% corresponded to San Simón sample, 31% to Cap Home, and 21% to Samil. This can be due to different causes (see discussion).

As the results were needed quickly, we decided not to work with individual reads and instead analyze the results of the co-assembly of the three samples. Since the coverage in all samples was low, the proportion of reads that could be assembled was low for all samples (26%, 24% and 23%), yielding just 599 contigs (but long ones: N50=35.5 Kb, longest contig, 179 Kb). To increase the information, we decided to use the option "--singletons" in SqueezeMeta, that takes all the unassembled reads and treats them as new contigs. In this way, all reads are represented in the analysis.

Finally, 64.228 contigs (N50: 6.600 bp) encoding for 256.774 ORFs were obtained. The analysis took approximately 4.5 hours to complete on our laptop. Therefore, the total length of the experiment was: Sampling: 4 hours. DNA extraction: 5 hours. Sequencing: 10 hours. Analysis: 4 hours, total 23.5 hours.

Inspection of the results in SQMtools and SQMxplore quickly determined that San Simón was the most interesting spot for sulfur metabolism, both in terms of abundance and presence of genes related to sulfur.

Different sulfur-related genes were found in the three different locations during the first day of sampling. Overall, the metagenome in San Simon Bay included a relatively higher abundance of sulfur genes (Figure 5). For example, SoxA (2.8.5.2) and SoxB (3.1.6.20) genes, thiosulfate sulfur transferases (2.8.1.1), TauACB and genes responsible for catabolizing sulfonamides (1.14.11.17) were relatively more abundant in San Simón Bay, suggesting an important presence of bacteria utilizing thiosulfate and bacteria incorporating taurine at this site.

**Figure 5:**
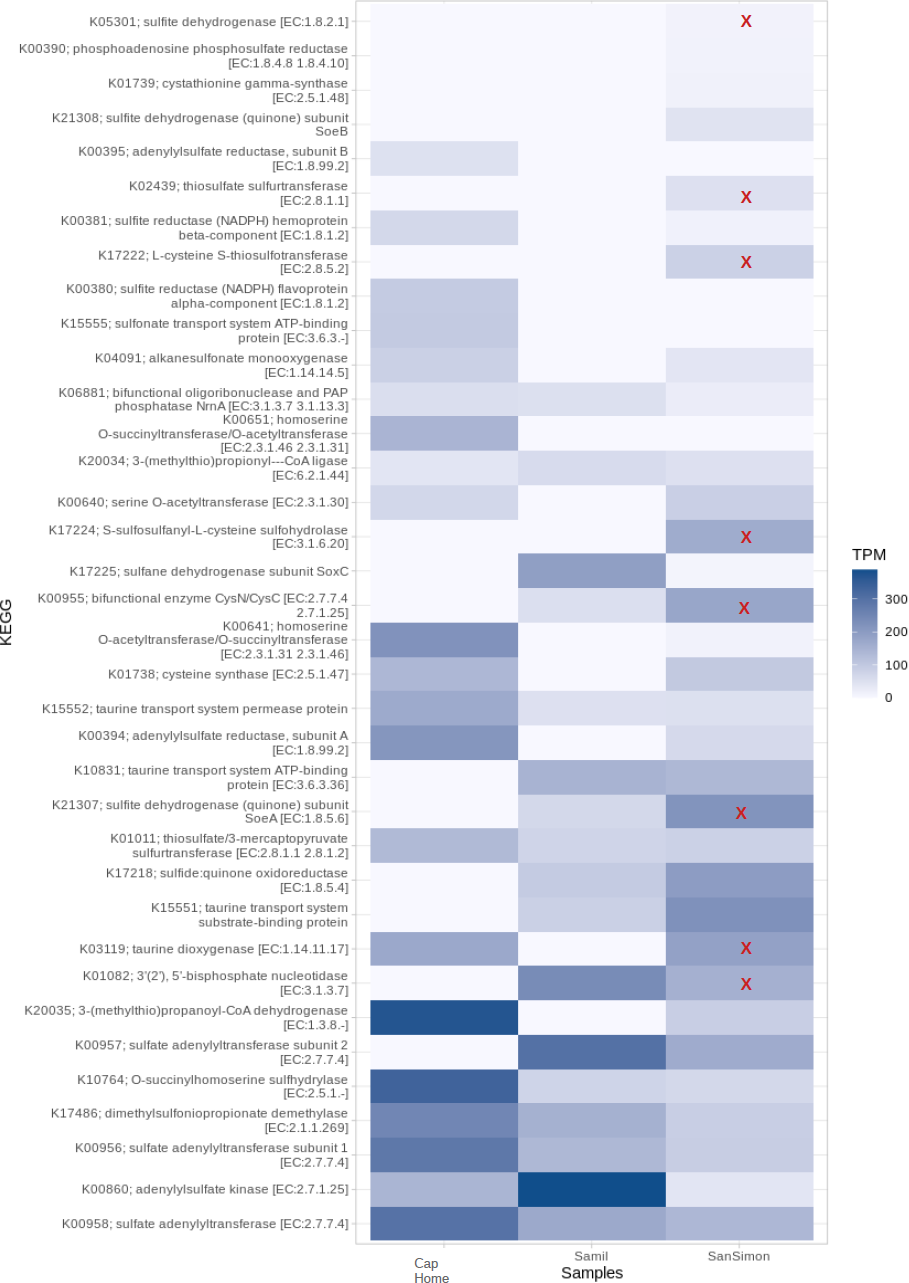
Relative abundances of sulfur genes in the three locations in Ria de Vigo. The rightmost column corresponds to San Simon sample. Genes driving the selection of this sampling spot for a second sequencing, as discused in the text, are marked. These are: SoxA (K17222, EC 2.8.5.2), SoxB (K217224, EC 3.1.6.20) genes, thiosulfate sulfur transferase (K02439, EC 2.8.1.1), Taurine dioxygenase (K03119, EC 1.14.11.17), dehydrogenation of sulfite (K21307, EC 1.8.5.6; K05301, EC 1.8.2.1) and sulfate reduction (K00955, EC 2.7.1.25,; K01082, EC3.1.3.7)

Similarly, dehydrogenation of sulfite (1.8.5.6, 1.8.2.1) and sulfate reduction (2.7.1.25, 3.1.3.7, CysND, CysH) were also relatively more abundant at San Simon Bay. Especially relevant was the presence of Sox genes, being the only sample in which we spotted the presence of SoxA and SoxB genes (Suppl Figure 3). Overall, the results from the first sampling day suggested that microbial communities from San Simon Bay will be of more interest for a second, more intensive (water column depth resolution) sampling.

This second sampling was done on July 14th 2022. We took two samples in San Simón sampling point, corresponding to two different depths (2 meters and 5 meters), so it was possible to characterize in detail sulfur metabolism of microbial communities from this station.

The concentration of extracted DNA was 29.7 ng/ml and 22.9 ng/ml for the samples at 2 and 5 meters, respectively. We performed sequencing during 10 hours using the same flow cell of the previous day. We aimed to obtain similar number of sequences for the two samples, therefore we adjusted concentrations to load the same amount of DNA for both. However, surprisingly, the total amount sequenced was 204 Mb and 32 Mb for both, emphasizing our difficulties to achieve equal sequencing depths (Table 1).

As time was not as demanding in this instance, a more complex approach was followed for the analysis, using co-assembly and the "doublepass" option of SqueezeMeta. This aims to discover extra genes by including an additional step of Blastx homology search on these parts of the sequences without gene prediction, or where the predicted ORF not matches anything in the nr database, pointing to a possible prediction mistake. The sample taken the previous day at the same location was also included in this analysis.

A summary of the results can be seen in Table 1. We obtained 1148 contigs in the assembly (Longest contig: 175796 bp) that contained approximately 30% of the reads. These were supplemented with 67186 singletons (unassembled reads). The final set of 68.334 sequences contained almost 500.000 ORFs, of which more than 400.000 matched some gene in the GenBank nr database (Benson et al, 2013).

During the second survey, interesting temporal and spatial (vertical) differences in sulfur-related genes in San Simon Bay metagenomes were found (Suppl Figure 4). Most of the sulfur-related genes found were relatively more abundant in surface samples (2 m) than close to bottom (5 m). This result may suggest, for example, that bacteria utilizing thiosulfate and bacteria incorporating taurine at this site are relatively more abundant in surface waters. On the other hand, a tendency to have higher relative abundance at surface waters on 14^th^ compared to 12^th^ July was found for some of the genes (e.g. SoxB, TauACB). These results suggest temporal changes in the relative importance of specific sulfur metabolisms in San Simón Bay.

Hence, the use of this in-situ strategy allowed to make an informed selection of the most interesting site at Ría de Vigo to perform an intensive metagenomic survey on sulfur-related genes, demonstrating the feasibility of this approach.

## Discussion

Analysis of metagenomic sequencing results is a work-intensive task involving several steps and different software tools, and requires careful statistical analysis to achieve the desired objectives (e.g. differences in functional or taxonomic diversity, or presence of particular genes or organisms). Therefore, bioinformatics expertise and powerful computational resources are needed.

To reduce this burden in resources and expertise, we have recently developed several software tools that provide a complete solution for all the bioinformatics involved in metagenomics. The SqueezeMeta software is a complete metagenomic pipeline that automatizes all steps of the analysis (Tamames & Puente-Sánchez, 2019a). It requires minimal user intervention, making it amenable to all kind of users, regardless of their bioinformatics expertise, and is able to work with limited computational resources, even allowing to analyze metagenomes on a laptop.

The second tool is the SQMTools software (Puente-Sánchez et al, 2020). This is a R library devoted to facilitate the statistical analysis of the results. The data generated by a SqueezeMeta run (e.g. contigs and gene sequences and annotations, aggregated functional and taxonomic profiles, and/or binning results) are loaded into a single R object, that can be explored with a set of simple functions allowing plot and chart drawing, performing multivariate analysis, or connecting to other popular analysis packages in microbial ecology. Nevertheless, the drawback was that users need to be somehow proficient in R usage to take full advantage of the power of this tool. To overcome this limitation, we have developed a third tool to facilitate the usage for all kind of users. This tool, named SQMxplore, includes a user interface for managing the data and allows sharing the results remotely with other users (Suppl Figure 5). SQMxplore is an application written using the R’s Shiny library that allows the loading of the tables created by SQMTools, as a result of a SqueezeMeta metagenomic analysis. This tool leverages the capacities of Shiny to provide an interactive graphical user interface, offering the possibility of visually inspect the tables, create and export customized plots, and perform multivariate analyses without the need of R programming. Shiny offers dynamic reloading of the results, so that any adjustments in the input data are immediately translated to the resulting tables or plots. It is also possible to upload the results to a web server, allowing remote users to interact with the data, thus facilitating considerably the discussion and dissemination of the results.

The combination of these three tools provides a complete solution for all the bioinformatic procedures involved in metagenomics, and together with the availability of portable sequencers, opens the way to be able to analyze metagenomes quickly and directly on the sampling spot. To test this capacity, we have sequenced and analyzed metagenomes from soils and marine waters.

We have shown that a portable laboratory fitting in a medium backpack can be enough to sequence and analyze a medium-size metagenome directly in the field. All devices are powered by batteries, thus not needing connection to a stable power source to work. Internet connectivity is not needed, unless the results wanted to be shared with remote users via the web interface provided by SQMxplore. Even in that case, the amount of data needed to be uploaded is tiny.

The weight of the portable laboratory is around 13 kg, so it can be carried by a single person for some time . This weight can be shared between different persons and/or put into some wheeled transporter if the terrain allows it. In the study of marine samples, the sample processing was performed in the base station to avoid carrying the bulky and heavy filtering devices. But if needed, these pieces of equipment could be added to the portable laboratory and powered with additional batteries. In this scenario, however, we have not tested yet if the ship movement, affecting the stability of the devices, can be an issue (Lim et al, 2014).

In laboratory tests devoted to prepare our next Antarctic campaign, we have found that the cold conditions severely affect the performance of the equipment, as observed by others (Gowers et al., 2019; Johnson et al., 2017). However, the usage of thermal insulated boxes filled with one or several battery-powered hand warmers, were enough to maintain moderately warm conditions that ensure the proper functioning of the instrumental.

When working with substrates like rocks, where microbial colonization is limited, we often face a problem related with the low concentration of DNA present in the samples. We ameliorated this drawback by processing higher amount of sample. In this study, it was necessary to process eight tubes with 200 ng of soil each, which were later collected in a single column, in order to concentrate as much as possible. Also, we realized that the setting of the bead beating procedure to lyse the cells was critical. We advise the usage of gentle conditions for this step. Vigorous beating could facilitate the breaking of the cells, especially if these are embedded in a solid matrix (Ammazzalorso et al, 2015), but it could also lead to extensive DNA fragmentation that would hamper the posterior sequencing. In terms of sequencing performance, it is much better to obtain fewer long sequences than many short ones, because the sequencing will be faster, consequently reducing the degradation of the flow cell. In addition, the preparation of the sequencing libraries is also conditioned by the size of the DNA fragments. Longer fragments will increase the ratio sequence/adapter, resulting in an excess of adapter. The different degree of DNA fragmentation will also hinder equalizing the contributions of different samples in multiplexing, because if one sample is more fragmented than the other(s), equal DNA concentrations can harbor different number of DNA molecules.

The long-term survival of flow cells is a real issue, especially when processing soil samples that are prone to have substances that can inactivate or damage the pores. After the initial sequencing runs, the number of available pores dramatically dropped, strongly hindering the reusing of the flow cells, and therefore increasing costs very much. In our experience, a cleaning/purifying previous step using magnetic beads to eliminate impurities improves the durability of flow cells, thus reducing the costs of in-situ sequencing.

Regarding the bioinformatic analysis, two different approaches for studying a metagenome can be used: to perform an assembly or co-assembly, or work with unassembled raw reads. The co-assembly provides a common reference for all the samples, making it easy the comparison, and generates longer sequences in the form of contigs more suited for the analysis, since they contain several genes that can increase the reliability of taxonomic and functional assignments. On the other hand, the lower is the amount of sequences, the less complete and comprehensive is the assembly. Using raw reads, thus skipping the assembly, has the advantage of using all information available, without discarding any reads. The main drawback is the more demanding computational costs, since this analysis is carried using Diamond Blastx (Buchfink et al., 2015), implying translation and homology searching of the six frames of each read.

To reach our goal of producing a full metagenomic analysis in less than 24 hours using a laptop as computing infrastructure, the analysis of raw reads is less feasible since it would take a longer time. Therefore, the co-assembly approach for analyzing the data was followed. Although the contigs obtained were rather long, only around 30% of the reads were assembled. To avoid discarding the unassembled reads, we used the singleton mode of

SqueezeMeta, which includes these as new contigs. The following steps of the analysis proceed as usual, with the prediction and annotation of putative ORFs. Gene predictors’ accuracy is reduced when the sequences are noisy, as it is frequent in minION sequencing, but this can be acceptable if we just want a glimpse at the functional profiles to, in our case, select the most interesting spot.

The previous strategy can be refined by using the “doublepass” option of SqueezeMeta when it is necessary to be more precise, such as during the second day of marine samples analysis. This mode includes a step in which the predicted ORFs are evaluated according to the results of homology searching. ORF showing a strong hit with high coverage are kept. An additional blastx search is performed in the parts of the sequence with discarded or no ORFs, including reliable hits as new ORFs.

In summary, we advise the following:

-Keep gentle conditions for the DNA extraction, especially when dealing with bead beating procedures. Extensive DNA fragmentation will hamper library preparation, reducing sequencing yield.
-Take into account that room temperature means 25°C. Performance of all reactions will degrade below that point. Take corrective measures such as the use of portable heaters.
-Put effort in purifying the extracted DNA. A contaminated DNA library can damage the flow cell very quickly.
-If the concentration is lower than the recommended 40 ng/ml, sequencing is possible but perhaps the ratio sequence/adapter may need be adjusted (add less adapter).
-A fast but representative analysis can be done by assemblying the sequences and adding unassembled reads to the resulting contigs (for this we use the – singletons option in SqueezeMeta).

We demonstrate here that it is possible to generate metagenomic information in less than one day, making it feasible to obtain taxonomic and functional profiles fastly and efficiently, even under field conditions. This capacity can be used in the future for real-time functional and taxonomic monitoring of microbial communities in remote areas.

## Acknowledgments

This research was funded by projects TRAITS (PID2019-110011RB-C31) and ROCKEATERS (PID2019-105469RB-C22) of Agencia Estatal de Investigación, Spanish National Plan for Scientific and Technical Research and Innovation. We thank the crew on the R/V Kraken and the ECIMAT team for their hospitality and professionalism during the cruises and lab work. We particularly thank professor Emilio Fernández for his advice and help during oceanographic cruises.

## Supplementary material

Suppl Figure 1: Functional profiles for sulfur metabolism from samples in La Palma island. Each box shows the abundance (in percentage) of the enzymes involved in this process. Boxes are splitted, with left half showing the abundance in the Teneguia sample, and the right half in San Antonio sample.

Suppl Figure 2: Comparison between the percentages of taxa and COG functions found in the two sequencing runs of MinION (in-situ) and Illumina (ex-situ).

Suppl Figure 3: Functional profiles for sulfur metabolism in Vigo Bay, first sampling

Suppl Figure 4: Functional profiles for sulfur metabolism in Vigo Bay, second sampling

Suppl Figure 5: Snapshot of the SQMxplore web tool, showing the analysis of he three samples in Ria de Vigo

## References

Ammazzalorso, A.D., Zolnik, C.P., Daniels, T.J., Kolokotronis, S. (2015). To beat or not to beat a tick: comparison of DNA extraction methods for ticks (*Ixodes scapularis*) PeerJ 3:e1147 https://doi.org/10.7717/peerj.1147

Benson D.A., Cavanaugh M., Clark K., Karsch-Mizrachi I., Lipman D.J., Ostell J., Sayers E.W. GenBank. Nucleic Acids Res. 2013 Jan;41(Database issue):D36-42. doi: 10.1093/nar/gks1195.

Breitwieser, F. P., & Salzberg, S. L. (2020). Pavian: Interactive analysis of metagenomics data for microbiome studies and pathogen identification. Bioinformatics 36(4), 1303–4. https://doi.org/10.1093/bioinformatics/btz715

Buchfink, B., Xie, C., & Huson, D. H. (2015). Fast and sensitive protein alignment using DIAMOND. Nature Methods, 12(1), 59–60. https://doi.org/10.1038/nmeth.3176

De Filippis, F., Parente, E., & Ercolini, D. (2017). Metagenomics insights into food fermentations. Microbial Biotechnology 10(1):91–102. https://doi.org/10.1111/1751-7915.12421

Deamer, D., Akeson, M., & Branton, D. (2016). Three decades of nanopore sequencing. Nature Biotechnology, 34, 518–524. https://doi.org/10.1038/nbt.3423

García-Gil, S. (2003). A natural laboratory for shallow gas: the Rías Baixas (NW Spain). Geo-Marine Letters, 23, 215–229. https://doi.org/10.1007/s00367-003-0159-5

Gowers, G. O. F., Vince, O., Charles, J. H., Klarenberg, I., Ellis, T., & Edwards, A. (2019). Entirely off-grid and solar-powered DNA sequencing of microbial communities during an ice cap traverse expedition. Genes 10(11):902. https://doi.org/10.3390/genes10110902

Johnson, S. S., Zaikova, E., Goerlitz, D. S., Bai, Y., & Tighe, S. W. (2017). Real-time DNA sequencing in the antarctic dry valleys using the Oxford nanopore sequencer. Journal of Biomolecular Techniques, 28(1), 2–7. https://doi.org/10.7171/jbt.17-2801-009

Koren S, Walenz BP, Berlin K, Miller JR, Bergman NH, Phillippy AM. (2017) Canu: scalable and accurate long-read assembly via adaptive *k*-mer weighting and repeat separation. Genome Res. 27(5):722–736. doi: 10.1101/gr.215087.116.

Kolmogorov M, Yuan J, Lin Y, Pevzner PA. (2019) Assembly of long, error-prone reads using repeat graphs. Nat Biotechnol. 37(5):540–546. doi: 10.1038/s41587-019-0072-8.

Laudadio I., Fulci V., Palone F., Stronati L., Cucchiara S., Carissimi C. (2018) Quantitative Assessment of Shotgun Metagenomics and 16S rDNA Amplicon Sequencing in the Study of Human Gut Microbiome. OMICS 22(4):248–254.http://doi.org/10.1089/omi.2018.0013

Li, D., Liu, C-M., Luo, R., Sadakane, K., and Lam, T-W., (2015) MEGAHIT: An ultra-fast single-node solution for large and complex metagenomics assembly via succinct de Bruijn graph. Bioinformatics, 31(10):1674–6. https://doi.org/10.1093/bioinformatics/btv033

Lim, Y. W., Cuevas, D. A., Silva, G. G. Z., Aguinaldo, K., Dinsdale, E. A., Haas, A. F., Hatay, M., Sanchez, S. E., Wegley-Kelly, L., Dutilh, B. E., Harkins, T. T., Lee, C. C., Tom, W., Sandin, S. A., Smith, J. E., Zgliczynski, B., Vermeij, M. J. A., Rohwer, F., & Edwards, R. A. (2014). Sequencing at sea: challenges and experiences in Ion Torrent PGM sequencing during the 2013 Southern Line Islands Research Expedition. PeerJ, 2, e520. https://doi.org/10.7717/peerj.520

Luo, W., & Brouwer, C. (2013). Pathview: An R/Bioconductor package for pathway-based data integration and visualization. Bioinformatics 29(14):1830–1. https://doi.org/10.1093/bioinformatics/btt285

Nogueira, E., Pérez, F. F., Rıos, A. F. (1997). Seasonal patterns and long-term trends in an estuarine upwelling ecosystem (Rıa de Vigo, NW Spain). Estuarine, Coastal and Shelf Science, 44(3), 285–300. https://doi.org/10.1006/ecss.1996.0119

Nowinski, B., Smith, C. B., Thomas, C. M., Esson, K., Marin, R., Preston, C. M., Birch, J. M., Scholin, C. A., Huntemann, M., Clum, A., Foster, B., Foster, B., Roux, S., Palaniappan, K., Varghese, N., Mukherjee, S., Reddy, T. B. K., Daum, C., Copeland, A., … Moran, M. A. (2019). Microbial metagenomes and metatranscriptomes during a coastal phytoplankton bloom. Scientific Data .6(1):129. https://doi.org/10.1038/s41597-019-0132-4

Puente-Sánchez, F., García-García, N., & Tamames, J. (2020). SQMtools: Automated processing and visual analysis of ’omics data with R and anvi’o. BMC Bioinformatics 21(1):358. https://doi.org/10.1186/s12859-020-03703-2

Quick, J., Ashton, P., Calus, S., Chatt, C., Gossain, S., Hawker, J., Nair, S., Neal, K., Nye, K., Peters, T., De Pinna, E., Robinson, E., Struthers, K., Webber, M., Catto, A., Dallman, T. J., Hawkey, P., & Loman, N. J. (2015). Rapid draft sequencing and real-time nanopore sequencing in a hospital outbreak of Salmonella. Genome Biology, 16(1), 114. https://doi.org/10.1186/s13059-015-0677-2

Quick, J., Loman, N. J., Duraffour, S., Simpson, J. T., Severi, E., Cowley, L., Bore, J. A., Koundouno, R., Dudas, G., Mikhail, A., Ouédraogo, N., Afrough, B., Bah, A., Baum, J. H. J., Becker-Ziaja, B., Boettcher, J. P., Cabeza-Cabrerizo, M., Camino-Sánchez, Á., Carter, L. L., … Carroll, M. W. (2016). Real-time, portable genome sequencing for Ebola surveillance. Nature, 530(7589), 228–232. https://doi.org/10.1038/nature16996

Ramírez-Pérez, A. M., De Blas, E., & García-Gil, S. (2015). Redox processes in pore water of anoxic sediments with shallow gas. Science of the Total Environment, 538, 317–326. https://doi.org/10.1016/j.scitotenv.2015.07.111

Tamames, J., & Puente-Sánchez, F. (2019a). SqueezeMeta, a highly portable, fully automatic metagenomic analysis pipeline. Frontiers in Microbiology 9:3349. https://doi.org/10.3389/fmicb.2018.03349

Tamames, J., Cobo-Simón, M. & Puente-Sánchez, F (2019b). Assessing the performance of different approaches for functional and taxonomic annotation of metagenomes. BMC Genomics 20, 960. https://doi.org/10.1186/s12864-019-6289-6

Turingan, R. S., Thomann, H. U., Zolotova, A., Tan, E., & Selden, R. F. (2013). Rapid Focused Sequencing: A Multiplexed Assay for Simultaneous Detection and Strain Typing of Bacillus anthracis, Francisella tularensis, and Yersinia pestis. PLoS ONE 8(2):e56093. https://doi.org/10.1371/journal.pone.0056093

Walsh, A.M., Crispie, F., Claesson, M.J., Cotter, P.D. (2017) Translating Omics to Food Microbiology. Annu Rev Food Sci Technol. 8:113–134. doi: 10.1146/annurev-food-030216-025729.

Zepeda, M. L., Sicheritz-Pontén, T., Gilbert, M. T. P. (2015). Environmental genes and genomes: understanding the differences and challenges in the approaches and software for their analyses, Briefings in Bioinformatics, (16), 5, 745–758. https://doi.org/10.1093/bib/bbv001

