## Supplementary figures and images for "In-situ metagenomics: A platform for rapid sequencing and analysis of metagenomes in less than one day"

### Supplementary Figures

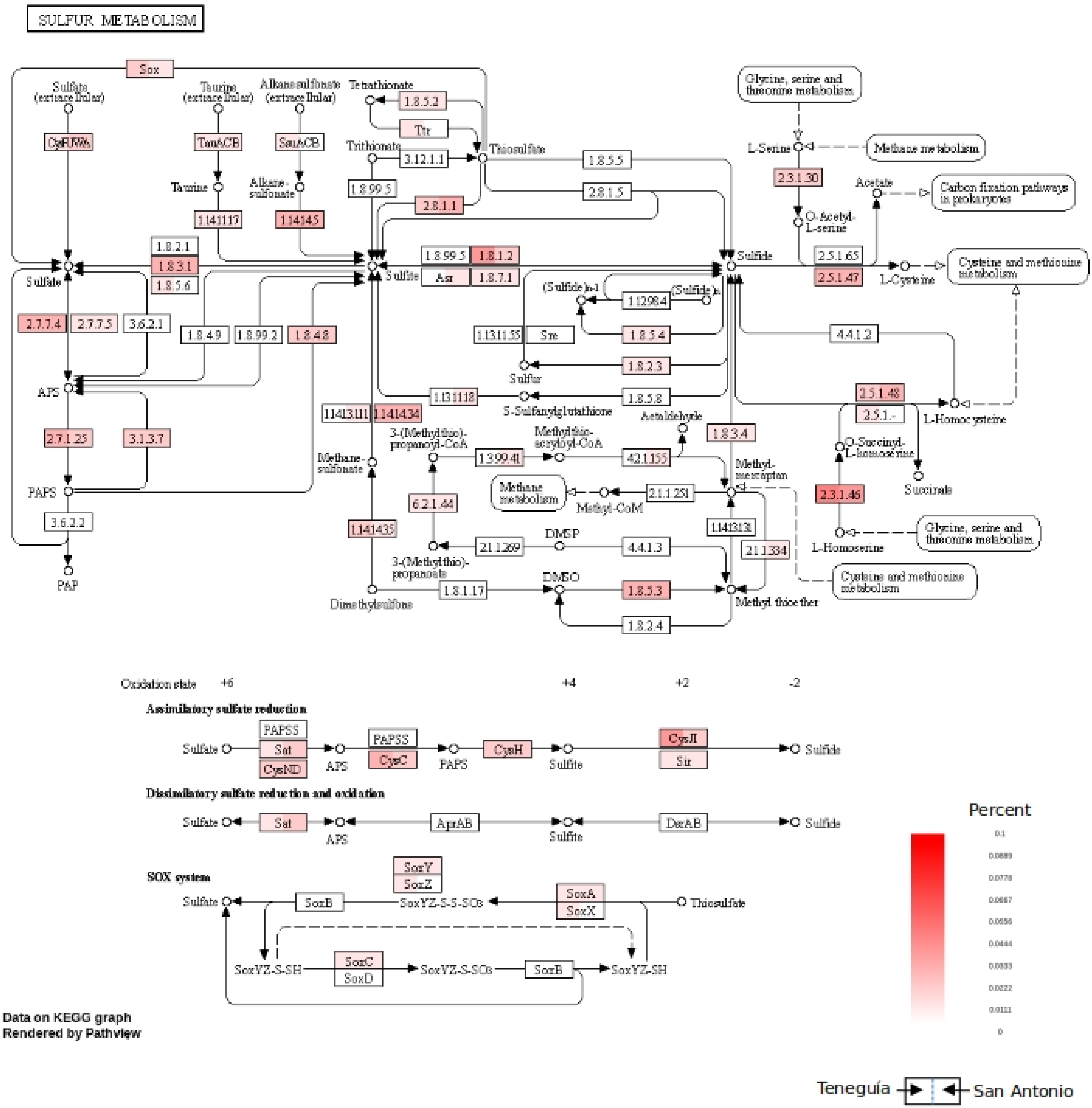


Suppl Figure 1

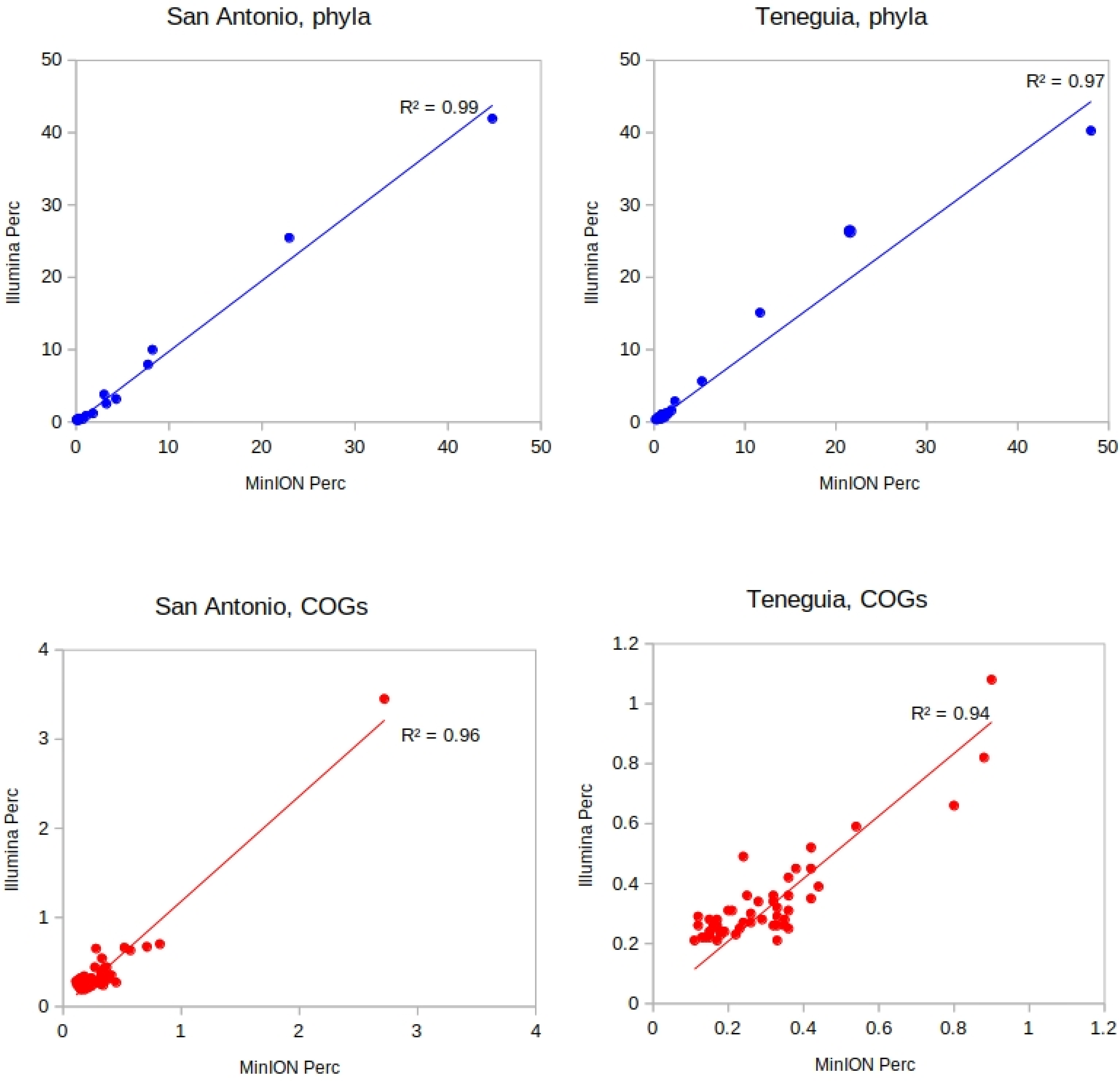


Suppl Figure 2

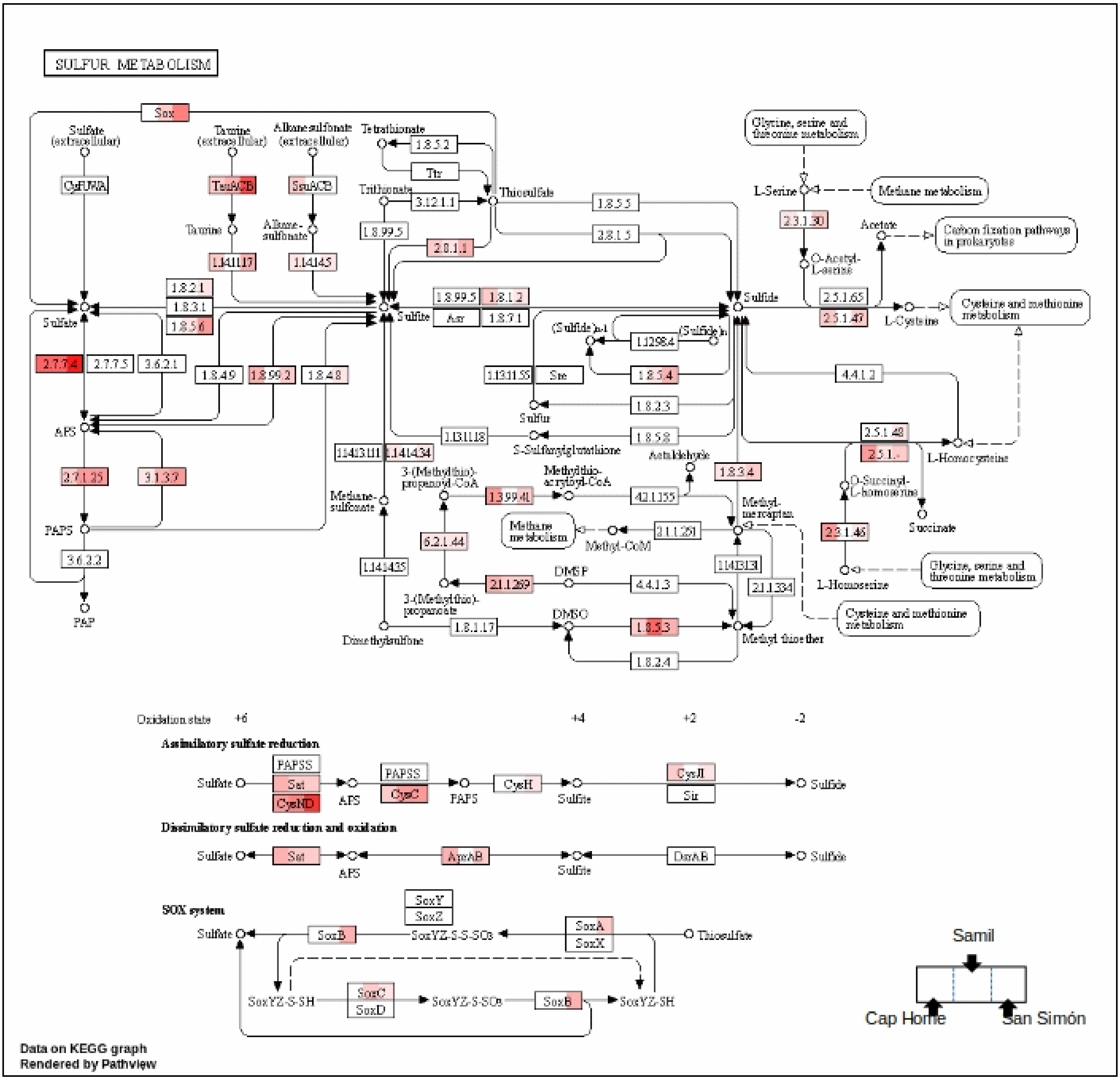


Suppl Figure 3

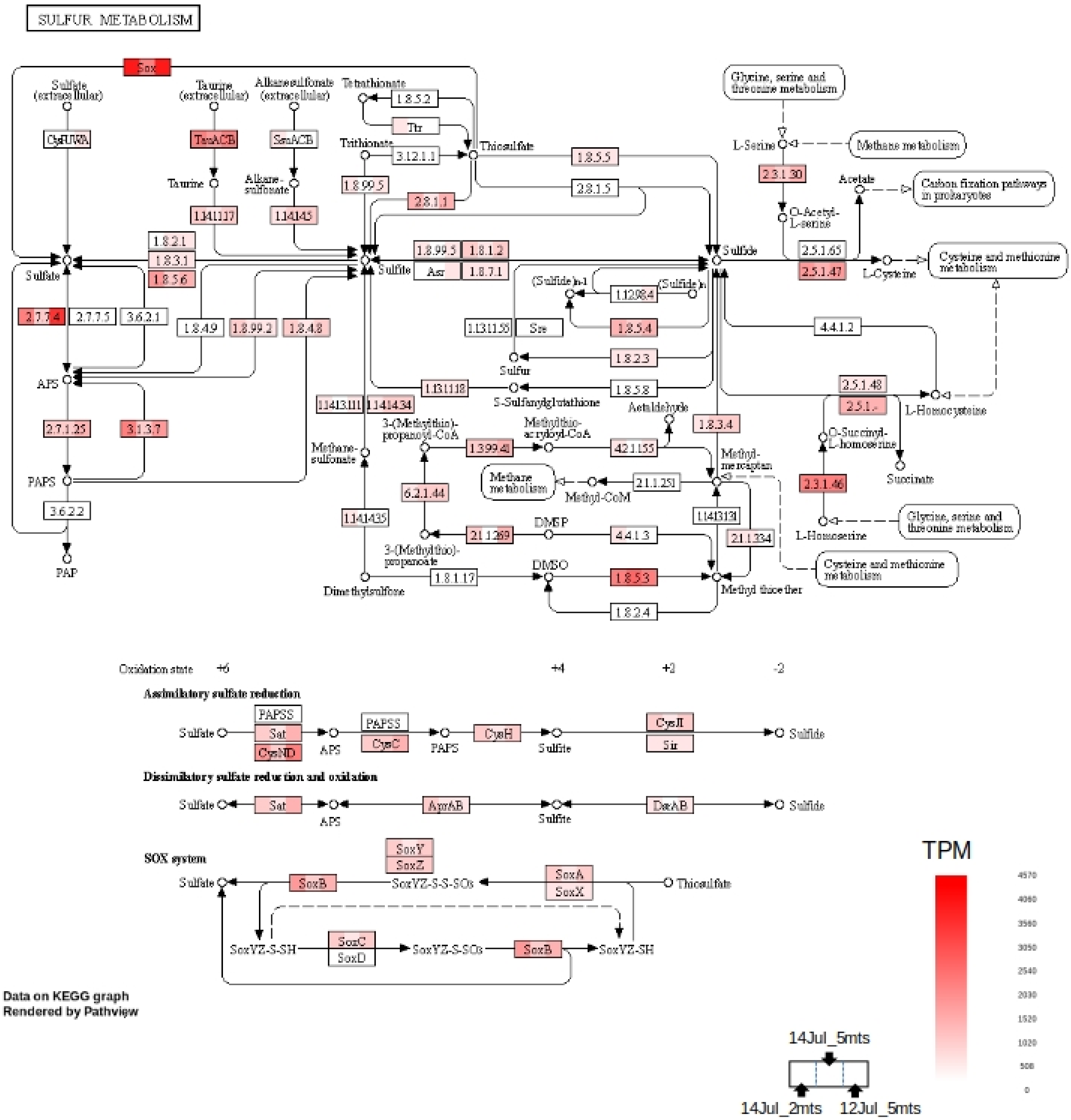


Suppl Figure 4

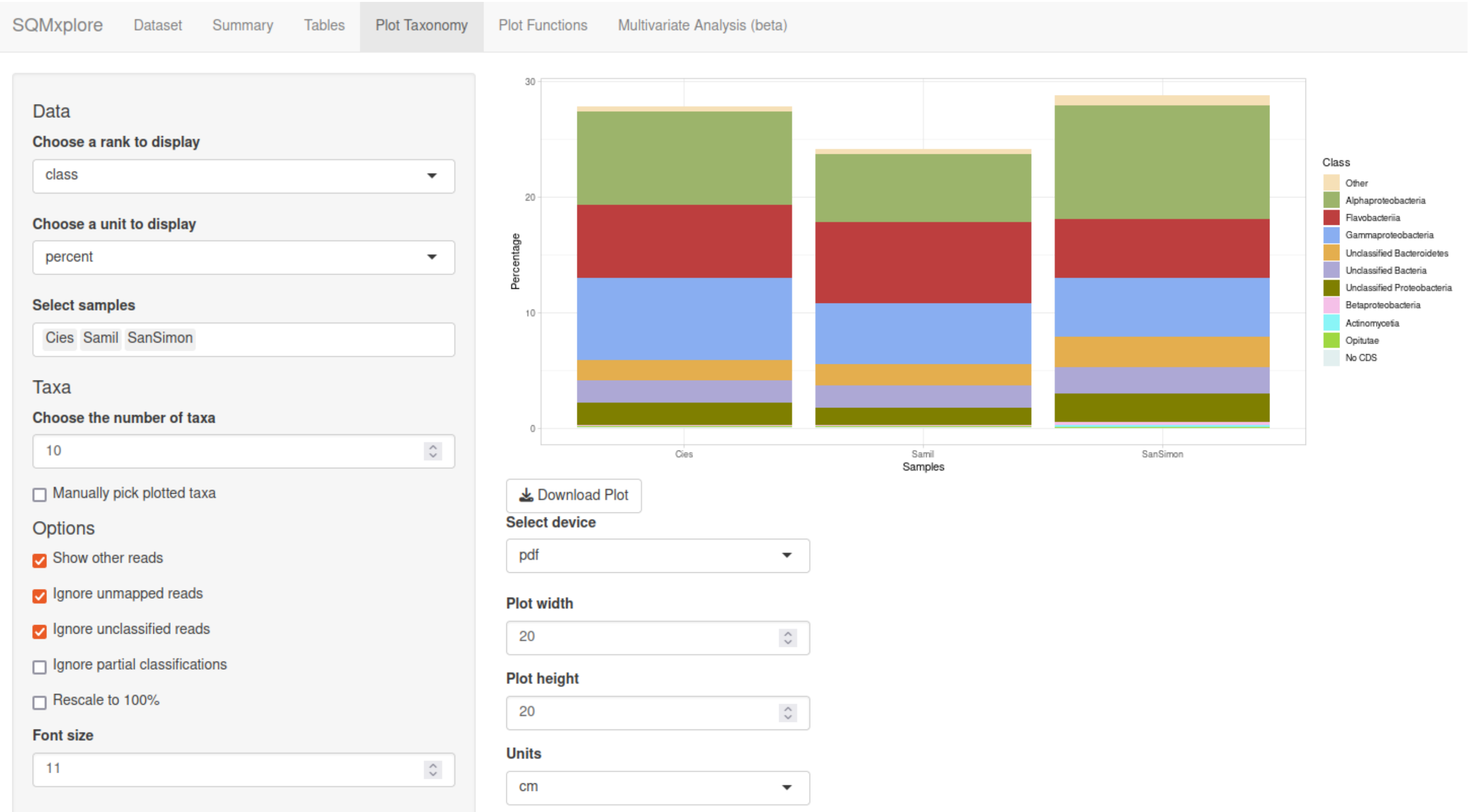


Suppl Figure 5
